# Location, location, location: Feeding site affects aphid performance by altering access and quality of nutrients

**DOI:** 10.1101/2020.08.10.244954

**Authors:** Vamsi J. Nalam, Jinlong Han, William Jacob Pitt, Shailesh Raj Acharya, Punya Nachappa

## Abstract

Feeding location on a plant can affect aphid reproduction and survival, however little is known about factors that influence aphid performance on specific plant parts. We determined performance and feeding behavior of soybean aphid (*Aphis glycines*) on stem, adaxial (upper) and abaxial (lower) leaf surfaces during early vegetative growth of soybean plants and analyzed the associated phloem sap composition. Stems harbored greater aphid populations and aphids had shorter development time on stems compared to adaxial and abaxial leaf surfaces. While aphids feeding on the stem took the longest time to begin probing, potentially due to higher density and length of trichomes, this did not impact aphid population growth. Once aphids began probing, the sieve elements were more conducive to feeding as evidenced by less salivation from the stem as compared to either leaf surface. Moreover, vascular sap-enriched exudates from stems had higher sugars and amino acids, which supported higher aphid populations in artificial diet feeding assays. The high quality of stems as a food source may in part explain the shorter development time and overall greater population of aphids observed on stems. In summary, our findings suggest that the choice of feeding location and performance of aphids on a specific plant is driven largely by accessibility to and the quality of nutrients rather than morphological factors.

## Introduction

Occurrence of insect herbivores on a given host plant is non-random and is affected by their preference for both plant phenology and physiology [1]. For phloem feeding insects such as aphids, the nutritional composition of the phloem sap is the main driver of fitness [2, 3]. The choice of feeding location can therefore affect aphid reproduction and survival by altering access and quality of food source [4, 5]. Several studies have demonstrated that within-plant variation in sugar and amino acid content of phloem sap parallels aphid feeding and performance [3, 6, 7]. For instance, many aphid species prefer young leaves with higher amino acid content over mature leaves with lower amino acid content [3, 8]. In fact, populations of the poplar leaf aphid (*Chaitophorous populicola* Thomas) appear to move from leaf to leaf on Eastern cottonwood, tracking the higher amino acid content in younger leaves [9]. Within individual plants, nutritional quality of leaves may vary depending on position and phenology [10-12]. As a consequence different aphid species prefer leaves of distinct age or growth stage indicating that aphids construct their own niche [2]. The cabbage aphid (*Brevicoryne brassicae* L.) exhibits higher fecundity and shorter generation time when restricted to feeding on flowers of canola (*Brassica napus* L.) compared to feeding on leaves. The cow pea aphid (*Aphis craccivora* Koch) demonstrates a definite preference for stem tissues compared to petioles and leaf blades, with significantly greater populations of aphids on stem tissues [13]. The choice of feeding location can also impact aphid behavior. For example, pea aphids (*Acyrthosiphon pisum* Harris) feeding on stems are more responsive to alarm pheromones and vibrations than aphids feeding on the abaxial or undersides of leaves [14]. Lastly, feeding location can alter vulnerability to natural enemies. Idris and Roff (15) found that the vertical and temporal distribution of the cotton/melon aphid (*Aphis gossypii* Glover) and its coccinellid predators is significantly greater in the lower stratum than in the middle and upper strata on different chili cultivars. These studies highlight the impact of feeding location on aphid biology through changes in feeding and oviposition/larviposition behavior.

The soybean aphid *Aphis glycines* Matsumura, a phloem-feeding insect native to Asia, was first detected in the U.S. in 2000, and is now present in all soybean growing regions in the United States [16]. The soybean aphid is considered the most economically important pest of soybean in the North Central United States. In addition to direct damage due to feeding, soybean aphids are competent vectors of many economically important plant viruses [16]. Under ideal conditions, soybean aphid populations can double in 1.5 days, and a single plant can harbor thousands of aphids [17]. A suite of Integrated Pest Management (IPM) strategies including, prophylactic neonicotinoid seed treatment, development of economic thresholds and injury levels, and deployment of aphid-resistant cultivars have been successful in controlling soybean aphids [18, 19]. However, the best option for control of soybean aphids remains scouting and foliar-applied insecticide, when the economic threshold of 250 aphids/plant is reached [20]. Hence, from a practical standpoint knowledge of within-plant distribution of aphids is important for accurate and rapid estimation of population densities. Within the plant, the distribution of soybean aphids varies with time. Early in the season, aphids are found on young leaves, higher in the canopy and as the season progresses aphids move to lower in the canopy in response to predation [8, 21]. A previous study reported that soybean aphids are most often found on the undersides of leaves [19]. However, there is no information on factors underlying soybean aphids’ preference for specific plant parts, which can have implications for scouting and treatment options.

The current study aimed to investigate how feeding location affects host plant acceptance and suitability for soybean aphids. In aphids, stylet penetration specifically phloem sieve element penetration is important for host plant acceptance and rejection [22]. However, only one study has reported on stylet penetration activities associated with aphid feeding at different locations on the plant. Prado and Tjallingii (23) did not find significant difference between stem and leaf surfaces, and neither between the leaf surfaces with respect to black bean aphid (*A. fabae* Scopoli) feeding on broad beans (*Vicia faba* L.); however, bird cherry-oat aphid (*Rhopalosiphum padi* L.) probed more on wheat stems and had more phloem ingestion on stems than on leaves. In the current study, we investigated aphid performance on stem, adaxial (upper) and abaxial (lower) leaf surfaces using life history assays and electrical penetration graph (EPG) analysis.Further, we examined plant morphology and nutritional composition as possible mechanisms underlying improved performance of aphids on a specific plant part. Our specific objectives were to: (1) determine the life history and population growth of aphids on adaxial, abaxial and stem surfaces, (2) determine feeding behavior of aphids on the three surfaces using the EPG technique, (3) determine if trichomes impacted aphid feeding, and (4) determine sugar and amino acid content from vascular sap-enriched stem and leaf petiole exudates of soybean and analyze their impact on aphid population growth.

## Materials and Methods

### Plant and insect maintenance

The soybean variety Asgrow AG3432 was used for all experiments. Plant growth and soybean aphid maintenance were as per Nachappa, Culkin (24). Briefly, plants were grown in Mastermix 830 soilless media (Mastermix, Quakertown, PA) in 15.24 cm diameter x 14.61 cm height plastic pots and watered three times per week. Plants were fertilized using Miracle Gro (Scott’s Company, Marysville, OH) solution once per week. The plants were maintained at 60-70% relative humidity, temperature of 24 ± 1 °C and a photoperiod of 16:8 (L:D) hours (h) at a photosynthetically active radiation (PAR) of 460 µmol/m^2^/sec in an environmental chamber. All experiments were initiated when plants were at the V1 stage or when the leaves of the first trifoliate were fully expanded. Soybean aphids (*A. glycines*) were originally collected from soybean fields at the Pinney Purdue Agricultural Center (PPAC), Wanatah, Indiana. In the laboratory, the aphid colony was maintained on AG3432 at temperature of 24 ± 1°C and a photoperiod of 16:8 (L:D) h in 30 × 30 × 76 cm insect cage (BioQuip, Rancho Dominguez, CA).

### Performance assay

Aphid population on adaxial, abaxial, and stem surfaces was evaluated by confining adult apterous aphids on the specific plant part using foam clip cages (36.5 × 25.4 × 9.5 mm) with no-thrips screen (BioQuip, Rancho Dominguez, CA). Ten adult apterous aphids were placed on either the adaxial, abaxial or stem surfaces and the total number of nymphs, apterous and alate adult aphids were recorded daily for four days, which is the time taken by soybean aphids to complete one generation under our laboratory conditions [25]. Soybean leaflets from the first trifoliates and stems between the first and second set of trifoliates were used for all assays. Each experiment had five replicates and the experiment was repeated three times (biological replications) for 15 plants total per treatment.

In separate experiments, life history traits of aphids including development time and longevity were assessed on whole plants. A single aphid was isolated on one of the three surfaces (adaxial, abaxial and stem) using clip cages. The adult was removed after 48 h and the development time or time to reach adulthood and offspring longevity were recorded. Adaxial and abaxial surface area measurements were identical, but limitations of the experimental setup created different surface areas for stems. However, in all setups there was ample room and resources for aphids to reproduce, feed, and survive, so we did not account for surface area in the analyses. There were 5 plants per treatment and the experiment was repeated five times (biological replications) for a total of 25 plants.

### Electrical penetration graph (EPG) analysis

Aphid feeding behavior on the abaxial, adaxial and stems of soybean plants was monitored using the electrical penetration graph (EPG) [26] technique on a GIGA 8 complete system (EPG Systems, Wageningen, Netherlands) [27]. The EPG experiments were performed on plants at the V1 stage or when the leaves of the first trifoliate were fully expanded. Adult soybean aphids were starved for 1 h prior to wiring. After wiring of aphids was completed, six plants, two per treatment were placed into a Faraday cage. The wired plant electrodes were then placed into the soil, and insect probes adjusted allowing for contact between the plant surface and the insect. Aphids were allowed to feed for eight hours, while the feeding behavior was recorded. Each feeding experiment was analyzed to determine the amount of time spent in each of the four main phases: pathway phase (PP), non-probing phase (NP), sieve element phase (SEP), and xylem phase (XP). Other parameters that were recorded include time to 1^st^ probe, time to first potential drop and the number of potential drops all of which provide an indication of aphid health [28]. EPG results were analyzed using Stylet+ software (EPG Systems, Wageningen, Netherlands). The experiment was repeated until a minimum of 25 replicates was obtained for each treatment (plant surface). There were 30 replicates for adaxial surface, 33 replicates for abaxial, and 25 replicates for stem surface. A behavioral kinetogram was constructed based on the number of transitional events for each waveform as per Ebert, Backus (29). The percent duration spent by the aphid in each waveform is represented by the area of the circles (see results). The arrows indicating transitions between the different waveforms were generated by dividing the transitional events for each waveform by the total number of transitional events.

### Trichome measurements

The number and length of trichomes on the first and second trifoliate leaves were determined using a digital camera connected to a Leica Zoom 2000 dissection microscope. The numbers of trichomes between and along the veins were counted on the image corresponding to 1 cm^2^ on the abaxial and adaxial leaf surface. For the stem sections, the number of trichomes were counted on the image corresponding to an area of 3.5 mm^2^. Trichome lengths were measured using Image J (https://imagej.nih.gov/ij/). Data presented are means from 18 samples (6 replications from three independent experiments).

### Collection of phloem and stem exudates

In separate experiments, plants (variety AG3432) were grown in pots with a 30.5 cm diameter (Myers Industries Marysville, OH, USA) until the V1 growth stage. Vascular sap-enriched stem and leaf petiole exudates were collected as per Nachappa et al [24]. In order to prevent bacterial contamination, trifoliates were cut at the base of the petioles or stems cut 2 cm above the soil surface were immersed in 50% ethanol, and then immediately moved to 0.05% bleach solution for no more than 2-3 seconds to achieve surface sterilization of the petiole and stem cut surfaces. The cut trifoliates and stems were then transferred to 1 mM EDTA solution (pH 8.0) until sampling was completed. Next, 1 cm above the previously cut surface for both the petioles and stems was cut, and three petioles with trifoliates and/or stems were weighed (to obtain fresh weight of sample) before being placed into the single well of a 6-well plate containing 6 mL of 1 mM EDTA. After completing transfer of all samples, the entire set-up (plant samples in 6-well plate) was placed in a terrarium with a clear lid lined with moistened paper towels for 24 h. At the end of the exudation period of 24 h, vascular sap-enriched stem and petiole exudates from three wells were then pooled per sample resulting in nine stems/petioles per pooled sample. Samples were then filtered through 0.2 µm pore size filters and lyophilized. After lyophilization, samples were eluted in 750 µL of 1mM EDTA solution and used in artificial feeding assays and for nutrient analysis (described below).

### Artificial feeding assay

An artificial diet developed for soybean aphids [30] was supplemented with lyophilized and reconstituted stem or leaf petiole exudates and used in artificial feeding assays. An artificial feeding chamber consisted of 55 mm petri dishes (VWR, Radnor, PA) with parafilm (Bemis, Neenah, WI) sachets containing 750 µL of diet and supplemented with 50 µL of leaf petiole exudate or 50 µL of stem exudates. Ten fourth instar aphid nymphs were placed in each feeding chamber and allowed to develop until adulthood. Total number of nymphs and adults were counted at the end of the experiment. The assay was conducted under laboratory conditions at ambient temperatures of 24 ± 1°C and 16:8 (L:D) h.

### Analysis of leaf petiole and stem exudates

Sucrose and amino acid content of the leaf petiole and stem exudates was quantified by gas chromatography-mass spectrometry analysis (GC-MS). The samples arranged in a randomized order were extracted and injected to GC-MS along with 3 QCs that were generated by combining a small aliquot of each sample. For quantification of sucrose, samples were first diluted by mixing 50 μL of sample with 450 μL of water. Then 50 μL of the diluted sample was mixed with 200 μL of 50% methanol in water containing 25 μg/mL of internal standard (D-Sucrose-13C12, 98%, Cambridge Isotope Laboratories, Inc.). After samples were well mixed, 30 μL were aliquoted and dried under nitrogen. For quantification of amino acids, samples were first diluted by mixing 80 μL of sample with 80 μL of 100% methanol. Then 3.2 µL of internal standard mix containing 200 µg/mL of L-^13^C_4_-asparagine (99%, Cambridge Isotope Laboratories, MA), ^13^C_6_,^15^N_2_-L-lysine (99 atom % ^15^N, 99 atom % ^13^C, 95% (CP), Sigma-Aldrich), threonine-^13^C_4_,^15^N (98 atom % ^13^C, 98 atom % ^15^N, Sigma-Aldrich), leucine-d_10_ (98 atom % D, Sigma-Aldrich), tryptophan-d3 (Santa Cruz Biotechnology) was added to each sample. After a thorough mixing, 100 μL were aliquoted, dried under nitrogen, and then stored at −80°C until analysis. The analysis of the dried samples was carried out as per Walsh, Miles (31). Briefly, the dried samples were resuspended in 50 μL of pyridine containing 25 mg/mL of methoxyamine hydrochloride and incubated at 60°C for 45 min. The samples were then vigorously vortexed for 30 s, sonicated for 10 min, and incubated for an additional 45 min at 60°C. Next, 50 μL of N-methyl-N-trimethylsilyltrifluoroacetamide with 1% trimethylchlorosilane (MSTFA + 1% TMCS, Thermo Scientific) were added, and samples were vigorously vortexed for 30 s, incubated at 60 °C for 30 min. Metabolites were detected using a Trace 1310 GC coupled to a Thermo ISQ mass spectrometer. Samples (1 μL) were injected at a 10:1 split ratio to a 30 m TG-5MS column (Thermo Scientific, 0.25 mm i.d., 0.25 μm film thickness) with a 1.2 mL/min helium gas flowrate. GC inlet was held at 285°C. The oven program started at 200°C for 30 s, followed by a ramp of 15°C/min to 330°C, and a 4 min hold. Masses between 50-650 m/z were scanned at 5 scans/sec under electron impact ionization. Transfer line and ion source were held at 300 and 260°C, respectively. Pooled QC samples were injected after every 6 actual samples.

### Statistical analysis

The aphid counts in the performance assays conformed to assumptions of ANOVA and no transformations were performed. The performance assay was designed as two-factor experiment (plant location and time) with repeated measures. Therefore, a repeated measure ANOVA was performed using the Fit General Linear Model function on Minitab 19. The development time and adult longevity data was not normally distributed; hence a non-parametric Kruskal-Wallis test was performed. A non-parametric Kruskal-Wallis test was also performed to compare the mean number and length of trichomes. The parameters measured for aphid feeding behavior data from the EPG experiments did not conform to the assumptions of ANOVA and therefore the data was subject to rank transformation following which ANOVA was used to determine the differences between groups followed by Tukey’s post-hoc test to determine differences between means that showed a *P-*value of ≤ 0.05. Two-sample t-test was performed to compare the mean quantities of sucrose and total free amino acids from the stem and leaf petiole exudates. All statistical analyses were performed using Minitab Version 19 (Minitab, Stat College, PA).

## Results

### Population growth of aphids

To determine performance of aphids on different plant parts, population of aphids, that is, total number of nymphs, apterous and alate adult aphids were measured daily. Aphid population significantly differed between feeding locations (F_2,168_=26.07, *P*<0.001), time (F_3,168_=35.56, *P*<0.001), but not the interaction between feeding location × time (F_6,168_=1.73, *P*=0.118). Soybean stems supported significantly greater population of aphids compared to either leaf surfaces on day 1 (F_2,44_=24.09, *P*<0.001) and this trend continued over the duration of the experiment (day 4, F_2,44_=13.07, *P* < 0.001) (Fig. 1). The development time or time to reach adulthood was significantly shorter on stems than on either adaxial or abaxial leaf surface (Table 1). Feeding location did not influence adult longevity; however, aphids on stems had increased longevity compared to both leaf surfaces albeit not statistically significant (Table 1).

**Table 1.** Life history parameters of soybean aphids reared on different plant parts.

|  | **Adaxial** |  | **Abaxial** |  | **Stem** |  | ***P*-value<sup>a</sup>** | ***H*-value<sup>b</sup>** |
| --- | --- | --- | --- | --- | --- | --- | --- | --- |
| Time to adulthood (days) | 8.1 ± 0.3 | a | 8.4 ± 0.3 | a | 6.9 ± 0.3 | b | **0.001** | 12.18 |
| Longevity (days) | 11.2 ± 0.8 |  | 12.0 ± 0.7 |  | 13.1 ± 0.4 |  | 0.208 | 10.14 |

<sup>a</sup>*P*-values that are significant are highlighted in bold. <sup>b</sup>*H*-value is the Kruskal-Wallis test statistic.

**Fig 1.**
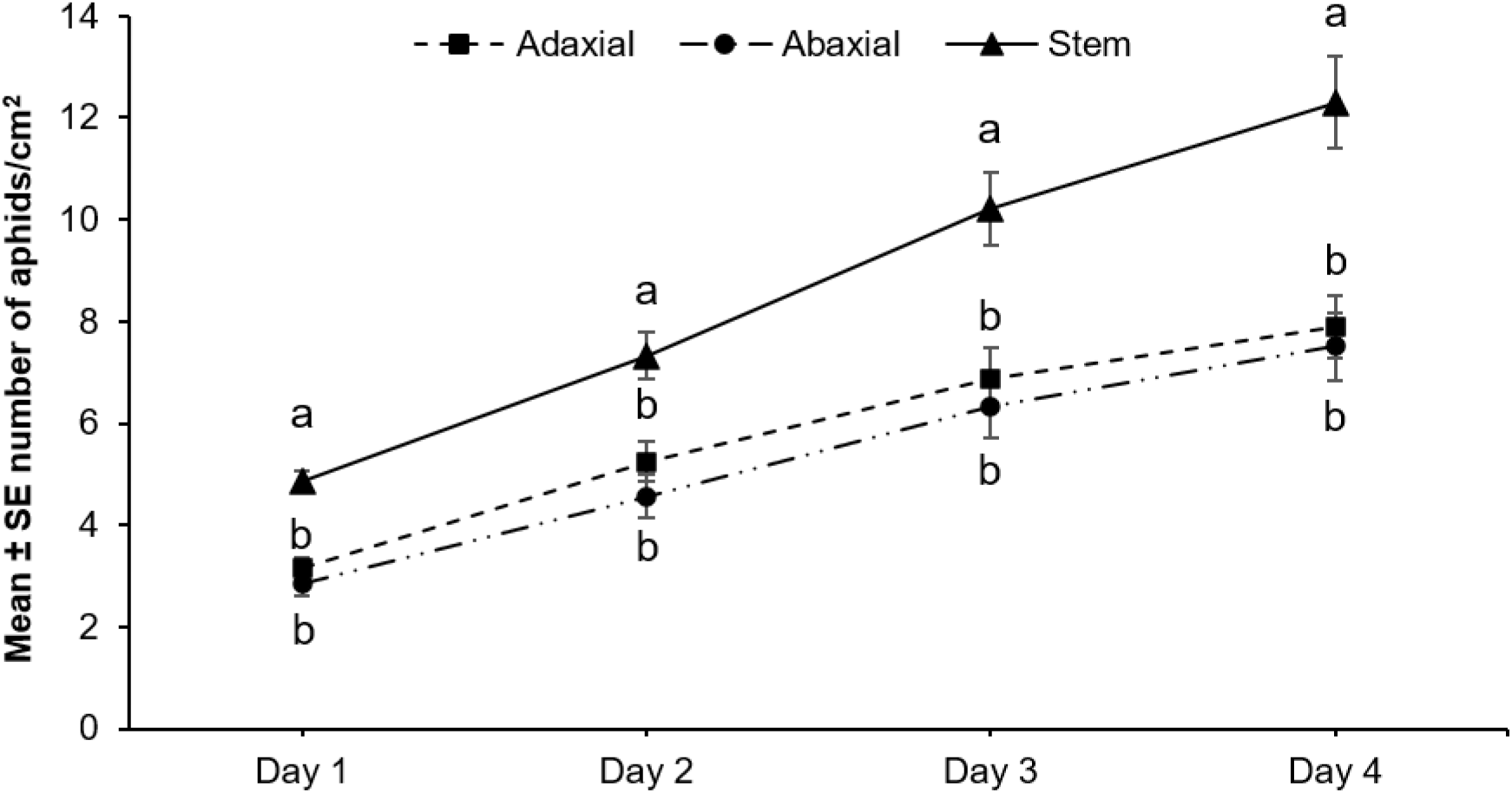
Population growth of aphids (total number of nymphs, apterous and alates) on adaxial, abaxial and stem surfaces. The numbers of adults and nymphs were recorded daily for four days. Five replications were performed for each host surface and the experiment was repeated three times. Different letters above the bars indicate values that are significantly different from each other (*P* < 0.05).

### Aphid feeding behaviors

We performed electrical penetration graph (EPG) analysis of aphid feeding behaviors on the three host surfaces. A behavioral kinetogram was constructed based on the transitions to and from each waveform and to represent the duration of time spent in each waveform (Fig. 2, Supplemental Table 1).

**Fig 2.**
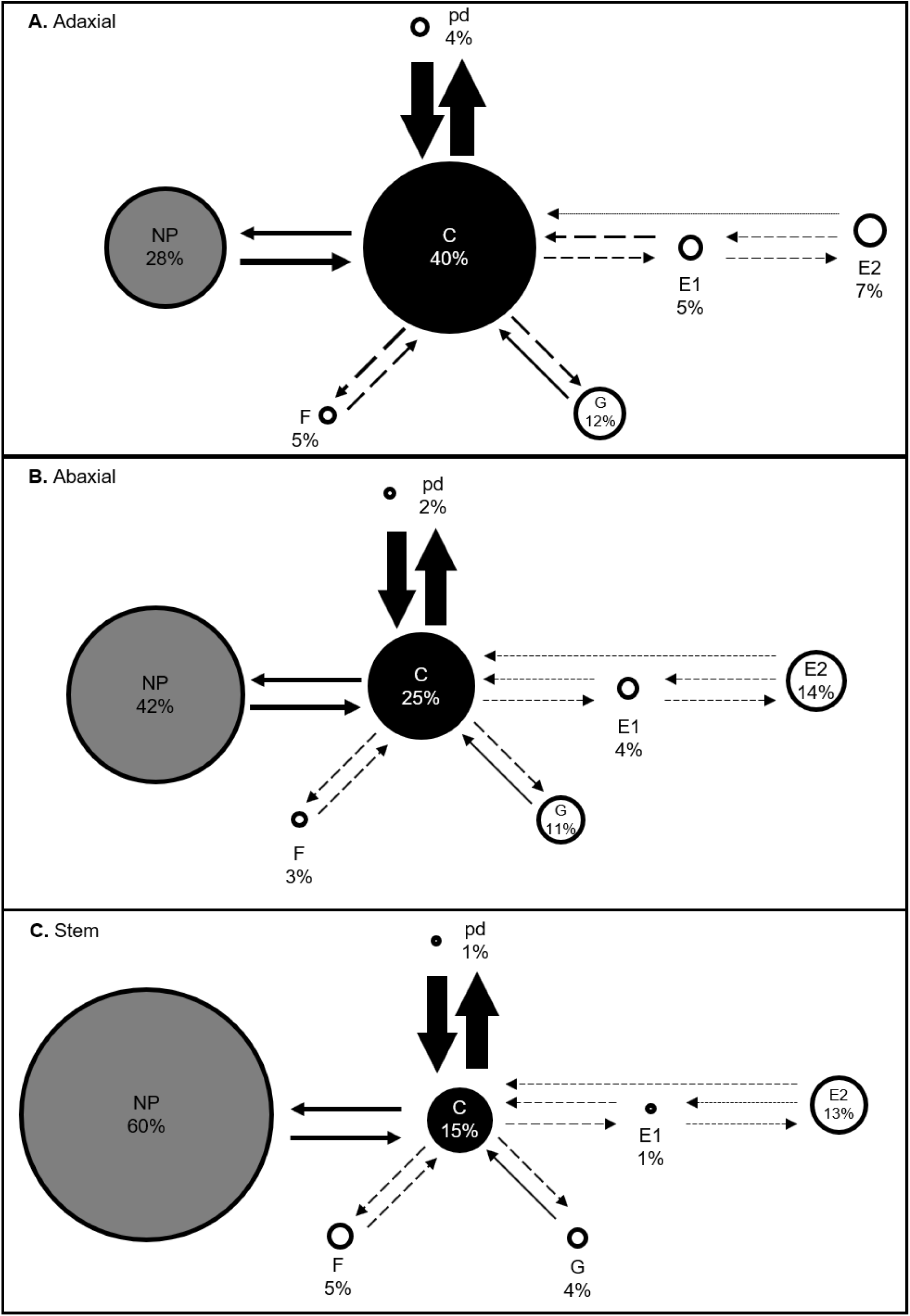
Behavioral kinetogram of aphid feeding behaviors. Circle areas represent the proportion of time spent by soybean aphids in different behaviors on the three different plant surfaces: (A) Adaxial, (B) Abaxial and (C) Stem. Arrows represent transitions with arrow thickness proportional to frequency. Values are averages of 30, 33 and 25 samples for the adaxial, abaxial and stem surfaces respectively. The time spent by the aphids in various activities (NP, non-probing phase; PP, pathway phase; SEP, sieve element phase; G, xylem phase) was analyzed by using rank transformation followed by GLM/ANOVA. The different colors of the circles indicate values that are significantly different (*P* < 0.05) from each other. Solid arrows represent values that are significantly different (*P* < 0.05) from each other and dashed arrows represent values that are not significantly different from each other.

#### Probing behavior

The time from the start of the EPG recording to the first probe varied significantly among aphids feeding on the three host surfaces (Table 2). Aphids feeding on the stem took the longest to begin probing (8.7 ± 2.5 min; Mean ± standard error of mean, SEM) whereas aphids feeding on the abaxial leaf surface took the shortest time to first probe (3.7 ± 1.7 min, Table 2). Aphids feeding on the stem also showed a significantly lower total number of probes and short probes (i.e. probes lasting less than 3 min) (Table 2) as compared to aphids feeding on either leaf surface. Although the number of probes by aphids feeding on all three host surfaces decreased during the course of EPG recording, aphids on the adaxial leaf surface displayed a higher amount of probing during the fourth and fifth hour of EPG recording as compared to aphids on the abaxial leaf surface or the stem (Supplemental Figure 1, *P* < 0.001).

**Table 2.** Non-phloem feeding behaviors of aphids feeding on adaxial, abaxial and stem surfaces of soybean plant.

| Parameters |  | Adaxial<br>(Upper)<sup>a</sup><br>N= 30 | Abaxial<br>(Lower)<sup>a</sup><br>N=33 | Stem<sup>a</sup><br>N=25 | P-value<sup>b</sup> | F-value (2,87) |
| --- | --- | --- | --- | --- | --- | --- |
| **Probing Behavior** |  |  |  |  |  |  |
| Time Spent in non-probing | (min) | 137.1 ± 22.6 a | 199.3 ± 27.7 ab | 288.2 ± 34.1 b | **0.003** | 6.33 |
| Time to 1<sup>st</sup> Probe | (min) | 7.5 ± 2.3 b | 3.7 ± 1.7 b | 8.7 ± 2.5 a | **0.005** | 5.35 |
| Number of probes |  | 29.1 ± 2.9 b | 23.4 ± 2.6 b | 12.0 ± 2.3 a | **<0.0001** | 14.63 |
| Number of short probes (C<3 min) |  | 18.9 ± 2.2 b | 15.7 ± 1.9 b | 8.2 ± 1.8 a | **<0.0001** | 12.47 |
| Total probing time | (min) | 342.8 ± 22.6 a | 280.6 ± 27.7 ab | 191.4 ± 34.2 b | **0.003** | 6.33 |
| Total duration of C | (min) | 210.3 ± 17.7 c | 128.8 ± 15.5 b | 75.0 ± 13.7 a | **<0.0001** | 16.35 |
| Number of potential drops (PD) |  | 136.0 ± 12.3 c | 80.5 ± 10.6 b | 35.7 ± 7.3 a | **<0.0001** | 21.43 |
| Average duration of PD | (min) | 0.14 ± 0.01 | 0.13 ± 0.01 | 0.25 ± 0.11 | 0.873 |  |
| Sum duration of PD | (min) | 17.6 ± 10.4 b | 10.9 ± 2.1 a | 17.7 ± 1.7 b | **<0.0001** | 11.94 |
| **Xylem Feeding** |  |  |  |  |  |  |
| Aphids with xylem phase | % | 73.3 (22/30) | 48.5 (16/33) | 28.0 (7/25) |  |  |
| Number of G |  | 11.4 ± 1.5 | 9.4 ± 1.3 | 7.3 ± 1.2 | 0.502 |  |
| Time to 1<sup>st</sup> G | (min) | 66.6 ± 20.8 | 62.7 ± 15.8 | 49.2 ± 8.2 | 0.715 |  |
| Mean duration of G | (min) | 11.4 ± 3.2 | 25.9 ± 9.5 | 39.1 ± 9.1 | 0.78 |  |
| Time spent in G | (min) | 75.5 ± 15.7 | 104.8 ± 15.3 | 82.5 ± 14.5 | 0.21 |  |

<sup>a</sup>Data presented are the means ± standard error of mean for aphids that displayed the behaviors.
<sup>b</sup>P-values that are significant are highlighted in bold.

#### Potential drops

Within the pathway phase (PP, also referred to as the probing phase, C), potential drops (pd) are found which represent intracellular punctures made by stylet tips [32]. These penetrations play an important role in host selection since they allow the aphid to sample cell contents. The total number of pd was lowest on stems followed by abaxial and adaxial leaf surfaces (Table 2). The total time spent by the aphids in pd was, however, lowest on the abaxial leaf surface. However, the time spent by the aphids in sampling cell contents i.e. average time spent in pd on the three host surfaces was not significantly different (Table 2).

#### Xylem Phase (G)

The active ingestion of xylem sap is an important mechanism for rehydration of aphids [33, 34]. There were no significant differences in time spent in G by aphids feeding on the three surfaces (Fig. 2A, B and C). However, a larger percentage of aphids (73.3%) feeding on the adaxial leaf surface entered the xylem phase compared to abaxial (48.5%) and stem (28.0%) surfaces (Table 2). Additional parameters related to the xylem phase of aphid feeding were evaluated including the number of G, the time to first G and the mean duration of G and no significant differences were observed for any of these parameters (Table 2).

#### Sieve Element Phase (SEP)

The SEP occurs when the stylet is located in the sieve element [35]. The SEP consists of the E1 phase during which salivation into the sieve element occurs and the E2 phase, correlated with passive ingestion of phloem sap along with concurrent salivation. Aphids can also display a single E1 phase which is not followed by E2 (phloem ingestion) which indicates the aphid’s inability to continuously feed from the sieve element and enter into E2. Among aphids feeding on the stem, only 60% displayed SEP compared to 80% and 82% on the adaxial and abaxial surfaces, respectively (Table 3). Considering only the aphids that did feed from the sieve element on the three host surfaces, a significant difference in total time spent in SEP was not observed (Fig. 2A, B and C, Table 3). A closer examination of aphid behavior in SEP shows significant differences for several parameters related to the E1 phase (Table 3). The number of E1 and single E1 (i.e. E1 waveforms without a subsequent E2 phase) were significantly lower in the stem and abaxial leaf surface as compared to the adaxial leaf surface (Table 3). Aphids on the stem also spent a significantly lower amount of time in the E1 phase (Table 3). The total duration of time spent in single E1 was the highest in the aphids feeding on the adaxial leaf surface and was significantly lower on the abaxial leaf surface and stem (Table 3). Additionally, aphids feeding on the stem exhibited reduced number of probes once they had successfully located and began fed from the sieve element (Table 3). With respect to the E2 phase (phloem ingestion), no significant differences were observed for any of the parameters measured. Although, not statistically significant, aphids did seem to spend longer time in E2 phase on stems compared to adaxial and abaxial surfaces (Table 3).

**Table 3.** Phloem feeding behaviors of aphids feeding on adaxial, abaxial and stem surfaces of soybean plant.

| *Sieve Element Phase* |  | **Adaxial**<br>**(Upper)<sup>a</sup>**<br>N= 30 | **Abaxial**<br>**(Lower)<sup>a</sup>**<br>N=33 | **Stem<sup>a</sup>**<br>N=25 | ***P*-**<br>**value<sup>b</sup>** | ***F*-value**<br>**(2,65)** |
| --- | --- | --- | --- | --- | --- | --- |
| Aphids with SEP | (%) | 80 (24/30) | 81.8 (27/33) | 60 (15/25) |  |  |
| Time in SEP | (min) | 73.0 ± 14.8 | 105.6 ± 25.4 | 112.8 ± 36.5 | 0.98 |  |
| Number of E1 waveforms |  | 9.0 ± 2.8 b | 3.9 ± 0.5 ab | 3.5 ± 1.4 a | **0.012** | 4.76 |
| Time to 1<sup>st</sup> E1 | (min) | 144.8 ± 18.8 | 125.4 ± 18.1 | 167.9 ± 32.8 | 0.206 |  |
| Total duration of E1 | (min) | 30.3 ± 5.6 b | 24.6 ± 8.7 ab | 8.3 ± 2.5 a | **0.014** | 4.54 |
| Number of single E1 |  | 5.6 ± 2.5 b | 1.5 ± 0.4 a | 1.6 ± 1.2 a | **0.002** | 6.74 |
| Total duration of single E1 | (min) | 12.6 ± 3.1 b | 6.5 ± 2.3 ab | 3.6 ± 2.4 a | **<0.0001** | 10.06 |
| Number of probes after 1st E |  | 13.5 ± 1.9 b | 11.3 ± 1.9 b | 2.2 ± 0.4 a | **<0.0001** | 8.61 |
| Total duration of E1 followed by E2 | (min) | 68.8 ± 14.6 | 97.3 ± 26.1 | 136.5 ± 38.5 | 0.99 |  |
| Number of E2 waveforms |  | 3.4 ± 0.5 | 2.3 ± 0.4 | 2.3 ± 0.3 | 0.42 |  |
| Total duration of E2 | (min) | 51.2 ± 13.7 | 80.9 ± 22.5 | 130.5 ± 38.3 | 0.092 |  |
| Time to 1<sup>st</sup> E2 | (min) | 199.4 ± 17.2 | 134.1 ± 17.3 | 198.8 ± 41.3 | 0.06 |  |

<sup>a</sup> Data presented are the means ± standard error of mean for aphids that displayed the behaviors.
<sup>b</sup> *P*-values that are significant are highlighted in bold.

### Trichome density

There were significant differences between the three host surfaces with respect to trichome density and length (Table 4). The density of trichomes on the stem was significantly higher than either leaf surface, with the stems having 8.4 times and 6.5 times the densities on the adaxial and abaxial surfaces respectively (F_2,47_=62.4, *P* < 0.001). The abaxial leaf surface had a significantly higher density of interveinal trichomes as compared to the adaxial surface (F_1,35_=22.5, *P* < 0.001). A significant difference was also observed with respect to trichome lengths, with trichomes on the stem measuring twice as long as trichomes from either leaf surface (Table 4). The density and length of trichomes along the veins was significantly higher on the abaxial leaf surface as compared to the adaxial leaf surface (F_2,83_=239.55, *P* < 0.001).

**Table 4.** Mean ± SE trichome densities on adaxial, abaxial and stem surfaces of soybean plant.

| Location of trichomes |  | Adaxial<sup>a</sup> | Abaxial<sup>a</sup> | Stem<sup>a</sup> |
| --- | --- | --- | --- | --- |
| *Interveinal* |  |  |  |  |
| Number of trichomes | (per mm<sup>2</sup>) | 3.29 ± 0.33 a | 4.24 ± 0.34 b | 27.60 ± 1.75 c |
| Length of trichome* | (mm) | 7.03 ± 0.37 a | 6.64 ± 0.42 a | 12.68 ± 0.62 b |
| *Veins* |  |  |  |  |
| Number of trichomes | (per mm) | 0.16 ± 0.01 a | 0.39 ± 0.01 b | n.a. |
| Length of trichome* | (mm) | 0.69 ± 0.03 a | 0.76 ± 0.02 b | n.a. |

<sup>a</sup> Within a row, values followed by same letter are not different ( $P < 0.05$ ).

### Sugars and amino acids

Given that aphids had higher population growth and shorter development time on stems, we sought to determine if differences in phloem sap composition between the stems and leaves contributed to the observations. Vascular sap-enriched stem and leaf petiole exudates analyzed using gas chromatography mass spectrometry (GC-MS), showed that the quantity of both sucrose and total free amino acids are significantly higher in the stem-exudates as compared to leaf petiole exudates (t = −3.42; *P*=0.009 and t = −2.15, *P*=0.008, respectively; Fig. 3A and B). Of the 18 amino acids measured, the concentration of 12 amino acids are significantly different (Table 5). Eight of the 12 amino acids, aspartic acid, tyrosine, asparagine, threonine, leucine, alanine, phenylalanine and proline are significantly higher in the stem. On the other hand, the concentrations of lysine, tryptophan, serine and glutamine are significantly lower in the stems as compared to the leaf petiole exudates (Table 5).

**Table 5.** Concentrations of individual amino acid (ng/gm fresh weight) in stem- and leaf petiole-exudates of soybean plants.

| **Amino Acids** | **Leaf petiole<sup>a</sup>** | **Stem<sup>a</sup>** | ***P*-value<sup>b</sup>** |
| --- | --- | --- | --- |
| Alanine | 0.12 ± 0.01 | 0.48 ± 0.1 | **0.004** |
| Arginine | 11.28 ± 1.94 | 20.49 ± 4.76 | 0.109 |
| Asparagine | 2.27 ± 0.31 | 8.08 ± 0.83 | **0.004** |
| Aspartic Acid | 38.08 ± 5.41 | 93.96 ± 28.84 | **0.016** |
| Glutamic Acid | 0.67 ± 0.04 | 0.61 ± 0.09 | 0.423 |
| Glutamine | 0.21 ± 0.01 | 0.04 ± 0.01 | **0.004** |
| Glycine | 0.3 ± 0.04 | 0.42 ± 0.09 | 0.262 |
| Isoleucine | 0.92 ± 0.02 | 0.83 ± 0.11 | 0.055 |
| Leucine | 0.7 ± 0.02 | 1.47 ± 0.18 | **0.004** |
| Lysine | 0.8 ± 0.08 | 0.43 ± 0.05 | **0.004** |
| Methionine | 2.12 ± 0.45 | 1.14 ± 0.27 | 0.201 |
| Phenylalanine | 0.12 ± 0.03 | 0.47 ± 0.16 | **0.006** |
| Proline | 0.03 ± 0.01 | 0.35 ± 0.08 | **0.004** |
| Serine | 0.22 ± 0.02 | 0.16 ± 0.02 | **0.037** |
| Threonine | 2.26 ± 0.02 | 3.54 ± 0.36 | **0.004** |
| Tryptophan | 3.87 ± 1.05 | 0.34 ± 0.08 | **0.004** |
| Tyrosine | 1.06 ± 0.29 | 8.33 ± 1.25 | **0.004** |
| Valine | 4.69 ± 0.15 | 5.85 ± 0.52 | 0.055 |

<sup>a</sup> Values represent mean ± standard error of n=6 replicates.
<sup>b</sup> *P*-values that are significant are highlighted in bold.

**Fig 3.**
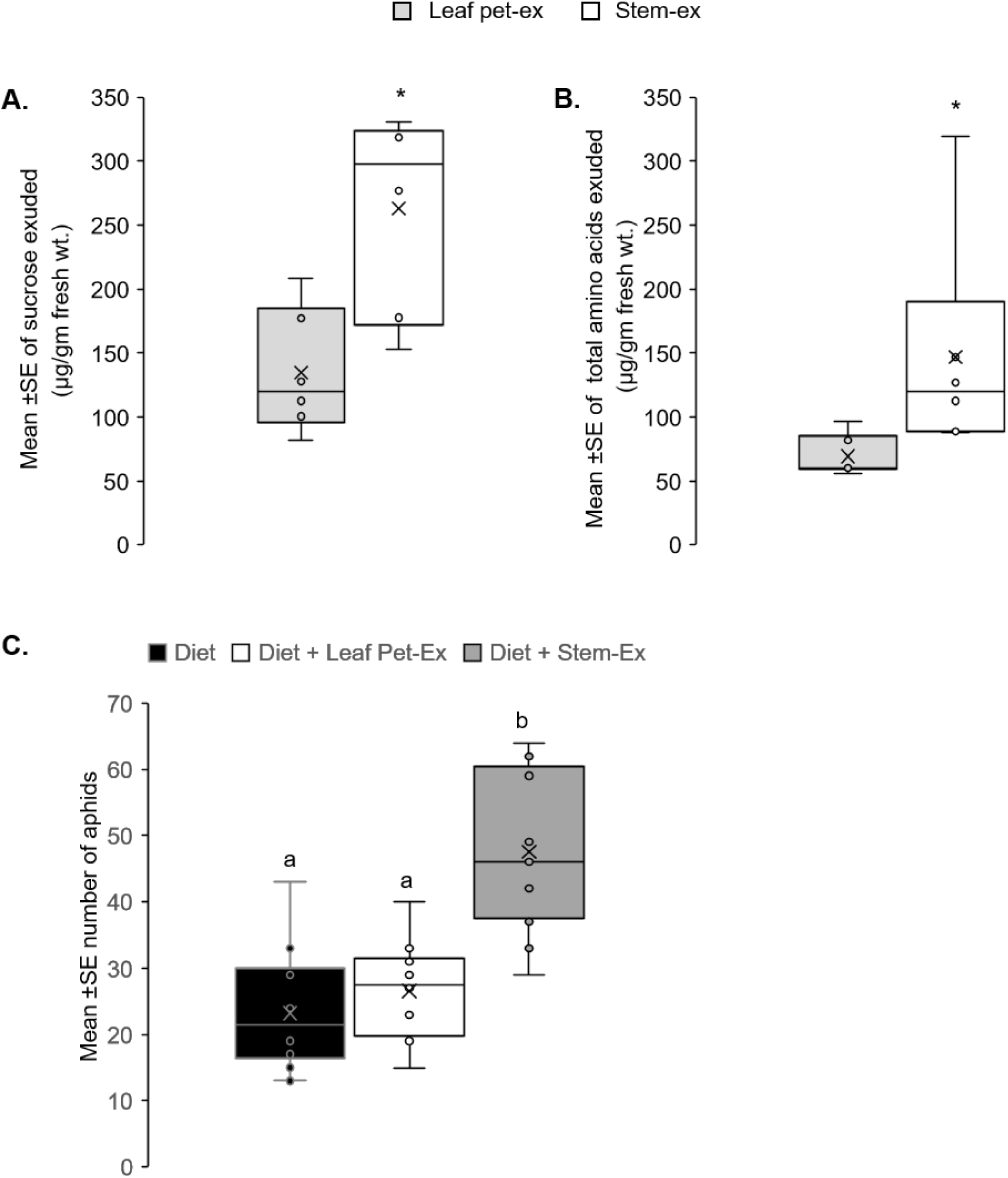
Phloem sap quality of stem supports higher aphid populations. GC-MS was used to determine the amounts of sucrose (A) and amino acids (B) in exudates collected from the stem and from leaf petiole exudates. Three biological replicates and two technical replicates were used to determine sucrose and amino acid quantity. Artificial diet assay with stem and petiole exudates (C). Comparison of aphid numbers on artificial diet supplemented with appropriate quantities of stem- and leaf petiole exudates. Control was artificial diet alone (diet). Aphid numbers were monitored for 4 days and data for day 4 is presented. The experiments were repeated 3 times with 4 replications each time. In (A) and (B) significant differences (*P* < 0.05) are indicated by asterisks (*). For (C), different letters above the bars indicate values that are significantly different (*P* < 0.05) from each other.

Artificial feeding assays were conducted using diet-enriched with either stem exudates, leaf petiole exudates or diet alone to determine the impact of phloem composition on aphid population growth. The population of aphids reared on stem exudate supplemented artificial diets was 2 times greater than artificial diet alone and 1.7 times greater than diet supplemented with leaf petiole exudates (F_2,41_=25.24, *P*<0.001; Fig. 3C). Taken together, these data suggest that stems represent a higher quality food source for aphids.

## Discussion

The choice of feeding location on a plant impacts insect fecundity and development directly through physiological mechanisms or indirectly through ecological mechanisms [15, 36-38]. In the current study, we show that soybean aphids have higher populations and shorter development time on stems compared to adaxial and abaxial leaf surfaces during early vegetative growth of soybean plants. While stems harbored a higher density and length of trichomes which resulted in aphids taking longer time to probe, no impact on aphid population growth was observed. Once the aphids began probing, the sieve elements were more accessible from the stem as evidenced by less salivation as compared to either leaf surface. The time spent in feeding from the sieve elements was greater in stem albeit not statistically significant. The quantity and quality of both sugars and amino acids was significantly higher in vascular sap-enriched exudates collected from the stems as compared to exudates collected from leaf petioles. The high quality of the stem as a food source may in part explain the shorter development time and overall high population of aphids observed on the stem. Additionally, in artificial diet assays, supplementing the diet with stem exudates supported higher populations of aphids. In summary, our findings suggest that the location of aphids on a plant is driven largely by accessibility and quality of nutrients rather than morphological factors.

Feeding location had a strong influence on aphid performance. In the current study, the population of soybean aphids (adults and nymphs) was significantly higher on stems compared to either leaf surface. Moreover, development time was significantly shorter, and the adult aphids lived longer on stems. These findings suggest that soybean aphids natural preference would be for stem, but Tilmon et al [39]. reported that soybean aphids are commonly found on underside of the leaves [19]. It is possible that aphid preference for specific plant parts varies over time throughout the phenology of the plant [40]. Aphid preference for stems has been observed in other aphid-host systems. Prado and Tjallingii (23) report that the stem is the preferred feeding site for bird-cherry oat aphids (*R. padi*) on wheat. Berberet, Giles (13) report similar behaviors for cowpea aphids (*A. craccivora*), which exhibited a strong preference for stems as opposed to petioles and leaf blades.

A limitation of our experimental design is that experiments were initiated when plants were at the V1 or vegetative stage, when the leaves of the first trifoliate were fully expanded. It is possible aphid preference for the stem occurs at the early vegetative stage of plant growth and changes in later vegetative and reproductive stages of the plants. Indeed, within-plant distribution of aphids is dynamic over the season [20]. Soybean aphids have greater rates of reproduction during vegetative stages than late reproductive stages presumably because of the lack of amino acids in the phloem when senescence occurs [40]. In contrast, more recent studies have shown that plant growth stage effects are not as pronounced [41-43]. Rutledge and O’Neil [41] found that life history traits of soybean aphids showed no difference on different growth stages of soybean plants. It is also important to note that predators [44] and abiotic factors such as high temperature and rainfall that may influence population growth and within-plant distribution of aphids. Nevertheless, mechanisms driving within-plant distribution of aphids needs further investigation. Future research should focus on soybean aphid preference and performance in both laboratory and field conditions and over longer temporal scales to elucidate any potential density dependent changes in preference and performance, in addition to identifying aphid performance during varying plant growth stages.

After initial plant contact, aphids use their stylets to probe the epidermis and this process is crucial for host plant and feeding site selection [23]. The first few stylet penetrations are usually confined to the epidermis and are usually brief (<3 min). These short probes are important indicators for gustatory cues as aphids ingest small amounts of plant cell contents to determine suitability. Our observation that soybean aphids show a significantly lesser number of short probes when feeding on stems suggests that non-vascular factors are involved [45, 46]. Trichomes on plants can act as morphological barriers interrupting and slowing down feeding by aphids [47-50]. On soybean, we found that there are significant differences in non-glandular trichomes between stem and leaf surfaces. Trichomes are denser and longer on stems compared to either leaf surface. This may have resulted in the reduced number of probes and longer time spent in non-probing observed on the stem. Longer non-probing periods have also been reported for bean aphids (*A. fabae*) and bird cherry-oat aphids (*R. padi*) feeding on stems [23]. However, in the current study trichomes did not negatively impact aphid populations, with stems supporting the highest populations. Similar to our observations, field studies indicate that trichomes do not affect the density of soybean aphids [51]. Additional evidence that trichomes do not impact soybean aphid populations comes from Pritchard (49), who used soybean isolines that differ in trichome density and found no impact on soybean aphid population growth. Although the two studies determined the impact of trichomes across different varieties or isolines, their findings confirm our observations within a single plant. It appears that soybean aphids have adapted to circumvent trichome-based plant defenses for their own advantage allowing them to possibly seek refuge from predators. Taken together, our results reveal that greater trichome density and length does not deter aphids from settling, feeding and reproducing on the stem compared to adaxial and abaxial surfaces.

The final behaviors observed during host plant and feeding site selection are phloem acceptance (salivation or first phloem phase, E1) and sustained phloem ingestion (E2 and sE2) [26, 52, 53]. After penetrating sieve elements, stylets secrete watery saliva into the phloem (E1 phase) in order to suppress phloem-based defenses and to allow for prolonged phloem sap ingestion (E2 phase). In aphids feeding on stems, we observed that the total number of E1 waveforms and the time spent in E1 was significantly lower on stems. Watery saliva contains several proteins, some of which have well-known biochemical activity that could either act as elicitors or suppressors of plant defense [54, 55]. The shorter salivation periods observed in aphids on stems suggests that plant defense responses and in particular phloem-based defenses in stems are reduced compared to leaf tissues. Interestingly, a recent study showed that concentrations of specialized metabolites was high in exudates of stems and yet stems supported higher aphid populations [2] suggesting that aphids can adapt to these compounds. Similar information regarding the presence of specialized metabolites in the phloem sap of soybean plants is not available, but it is likely the case for soybean aphids as well. Additional evidence for the hypothesis that defenses in stems is reduced comes from the observation that the number of probes after first phloem salivation is shortest on stems. In other words, aphids face fewer interruptions, in terms of host defense responses after they successfully locate and begin feeding. However, additional experiments focused on quantifying host defense responses in the stems versus leaves is required.

Phloem sap quality can differ within a plant which can affect aphid performance [2, 3]. Studies have linked population growth of soybean aphids with increase in nitrogen concentration in phloem [57, 58]. In accordance with this hypothesis, we found that the concentration of both sucrose and free amino acids were significantly higher in vascular sap-enriched exudates collected from the stems compared to exudates collected from leaf petioles. Moreover, certain amino acids such as asparagine, threonine, leucine, alanine, phenylalanine and proline were significantly higher in stem exudates. Asparagine is critical for soybean aphid development and fecundity. In artificial diet assays, lower than normal levels of asparagine cause longer development times and lower fecundity with significantly fewer soybean aphids maturing to adulthood [30]. Conversely, green peach aphids (*Myzus persicae*) reared on diets supplemented with asparagine display enhanced growth [59]. Alanine, leucine, and glutamic acid accounted for 43% of variations in the intrinsic rate of increase in populations of the green peach aphids (*M. persicae*) and cabbage aphid (*B. brassicae*) [60]. In artificial feeding assays, population of aphids reared on stem exudates was 2 times greater than artificial diet alone and 1.7 times greater than diet supplemented with leaf petiole exudates mirroring the results observed in whole plant assays.

In summary, our results demonstrate that the aphid performances on different plant parts is linked to accessibility and quality of phloem sap. Future transcriptomic and metabolomic studies focused on the sieve element and phloem sap may help to understand global changes in phloem chemistry to help address mechanisms underlining aphid performance on different plant parts. Overall, our findings provide insights into how plant physiology affects within-plant aphid population growth and these results may be useful in developing management plans for soybean aphids.

## Acknowledgements

This work was partially supported by the Office of the Vice President for Funding at Colorado State University to P.N and V.J.N, and by NIFA-AFRI Seed Grant 2017-08469 to V.J.N.

## Author Contributions

V.J.N and P.N. conceived and designed the research. J.H. and S.R.A maintained the aphid colonies. J.H., W.J.P. and S.R.A conducted experiments and analyzed data. V.J.N. and P.N. analyzed the data and wrote the manuscript. All authors read and approved the manuscript.

---

**S1 Fig. Aphid feeding behaviors vary in response to host surface**. Electrical penetration graph (EPG) was utilized to determine probing behavior on the adaxial, abaxial leaf surfaces and the stem in 8 h of recording time. Values are averages of 30, 33 and 25 samples for the adaxial, abaxial and stem surfaces, respectively. Error bars represent standard error of mean (SEM).

