## Supplemental Table 1 for "Location, location, location: Feeding site affects aphid performance by altering access and quality of nutrients"

**Supporting information**

| **Transitions** | **Adaxial** | **Abaxial** | **Stem** | ***P-value*** | **F-value^a^** |
| --- | --- | --- | --- | --- | --- |
| Total transitions | 366.2 ± 30.1 | 219.8 ± 26.5 | 111.0 ± 19.3 | **0.000** | 21.86_2,87_ |
| NP to C | 28.6 ± 2.9 | 23.1 ± 2.6 | 11.9 ± 2.3 | **0.000** | 13.94_2,87_ |
| C to NP | 23.2 ± 2.3 | 18.5 ± 2.0 | 9.3 ± 1.8 | **0.000** | 15.39_2,87_ |
| C to pd | 139.8 ± 12.5 | 80.4 ± 10.7 | 46.0 ± 8.1 | **0.000** | 21.73_2,87_ |
| pd to C | 127.5 ± 11.9 | 72.8 ± 9.7 | 43.3 ± 7.9 | **0.000** | 20.78_2,87_ |
| C to E1 | 6.7 ± 2.7 | 3.0 ± 0.5 | 3.1 ± 1.7 | 0.105 | 2.35_2,55_ |
| E1 to C | 6.3 ± 2.5 | 2.5 ± 0.5 | 5.3 ± 4.3 | 0.417 | 0.89_2,41_ |
| E1 to E2 | 3.4 ± 0.6 | 3.1 ± 0.4 | 2.3 ± 0.4 | 0.420 | 0.88_2,51_ |
| E2 to E1 | 3.0 ± 0.6 | 3.1 ± 0.4 | 2.3 ± 0.4 | 0.191 | 1.69_2,87_ |
| E2 to C | 1.6 ± 0.2 | 1.9 ± 0.3 | 1.3 ± 0.2 | 0.625 | 0.48_2,31_ |
| C to G | 9.9 ± 1.4 | 6.6 ± 1.1 | 5.8 ± 1.4 | 0.127 | 2.14_2,64_ |
| G to C | 9.2 ± 1.3 | 5.8 ± 1.1 | 4.3 ± 1.2 | **0.011** | 4.90_2,62_ |
| C to F | 9.7 ± 1.7 | 5.9 ± 1.3 | 7.0 ± 2.1 | 0.104 | 2.36_2,58_ |
| F to C | 8.5 ± 1.5 | 5.7 ± 1.2 | 5.4 ± 1.2 | 0.165 | 1.86_2,60_ |

**S1 Table. Transitional events observed for each waveform in aphids feeding from the three host surfaces.**

^a^*P*-values that are significant are highlighted in bold.
