## Supplementary figures and images for "Location, location, location: Feeding site affects aphid performance by altering access and quality of nutrients"

### Supplemental Fig 1

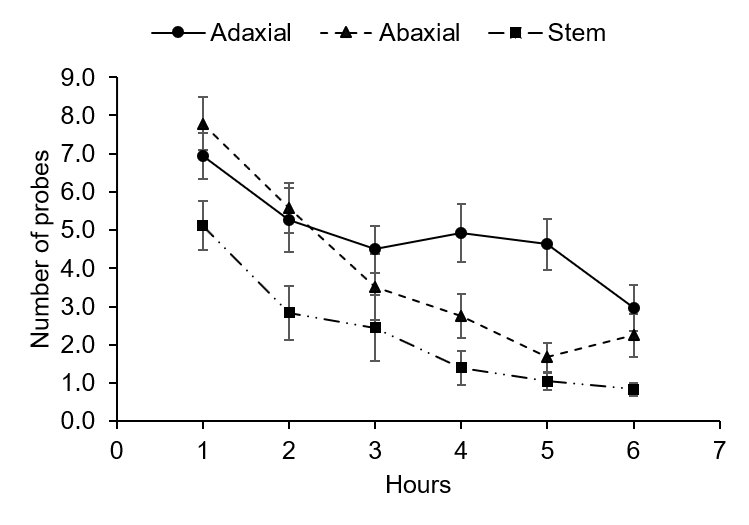
